# Dispersal behaviour as the outcome and trigger of multilevel selection in a social spider

**DOI:** 10.1101/866038

**Authors:** Zsóka Vásárhelyi, Jonathan N. Pruitt, István Scheuring

## Abstract

The facultatively social spider *Anelosimus studiosus* offers a unique opportunity for understanding how multilevel selection acts in natural populations. However, the importance of previous empirical studies are shaded by a conceptual debate about whether colony-level selection is truly present in these populations or not. Here we introduce a detailed individual based model, where practically all assumptions are supported by empirical data. The only element of the female *A. studiosus* life cycle missing from the literature is how maturing female spiders decide whether to disperse. This behavioural component we estimate with evolutionary simulations. This model is able to recapitulate the characteristic size and composition distributions of natural populations in different environments. The evolutionary simulations revealed that the optimal dispersal behaviour of a maturing female varies both with her ecological environment and behavioural phenotype. This finding is open for straightforward empirical testing. In agreement with empirical findings we have established parameter ranges where the population is prone to extinction without multiple-female nests. We propose that the dispersal behaviour of individuals is both the result and the prerequisite of multilevel selection in this species.

## Introduction

There are few debates more heated in evolutionary biology than that of the levels of selection (Nowak et al. 2010, 2011, Abbot et al. 2011, Kramer and Meunier 2016). Although the corresponding literature is still expanding rapidly, the tone of the debate has changed surprisingly little since the sixties (Lewontin 1970, Okasha 2006). Only recently, the concept of multilevel selection seemed to loosen this tension (Okasha 2006, Bijma and Wade 2008, Eldakar et al. 2010, Fisher et al. 2017, Fisher and Pruitt 2019), yet whenever the term *group selection* appears in print, a new wave of criticism and conceptual dispute arises. This was the case when Pruitt and Goodnight (2014) published their results claiming that the facultatively social spider *Anelosimus studiosus* shows signs of undergoing strong group selection (Grinsted et al. 2015, Gardner 2015, Pruitt and Goodnight 2015).

Most *A. studiosus* live subsocially (Brach 1977, Furey 1998), where mothers provide extended care for their offspring but latent aggression causes juveniles to disperse from their natal web. However, in the northern and coldest parts of its range, subadults are not driven away but instead remain in the natal web forming multigenerational multifemale colonies (Furey 1998, Jones et al. 2007). In such social populations, *A. studiosus* shows two distinct behavioural phenotypes, *aggressive* or *docile*. These two behavioural types specialise on different tasks in mixed colonies (Riechert and Jones 2008, Pruitt et al. 2008, Wright et al. 2014). Pruitt and Goodnight (2014) demonstrated that in environments of different resource levels, social *A. studiosus* colonies show opposing trends in their optimal size-composition relationships. That is, in one environment larger the colonies, larger the proportion of aggressives, and the opposite in the other environment. These opposing trends are partly the result of the differential extinction of colonies due to between-site variation in the factors causing colony extinctions. Experimental colonies placed in foreign environments adjust their compositions in a manner that tracks the optimal size-composition of their site-of-origins, even generations later. These data led the authors to conclude that selection acts primarily on groups and resulted in an adaptation in a colony level trait.

Their results, as expected, were heavily critiqued. Both initial responses to their paper highlighted the need to simultaneously account for individual-level selection in the analysis (Grinsted et al. 2015, Gardner 2015). Furthermore, Gardner (2015) went as far as to suggest not to invoke group selection until it is proved that within- and between group interests conflict.

In response, Pruitt et al. (2017) performed several follow-up experiments showing that within- and between-group interests do not conflict in this species, mostly because both individual fitness and group success is primarily driven by rapid group extinctions. They conclude, however, that the tendency to optimise group composition still seems to be a group-level adaptation by their definition, because it is transmitted down colony lineages in a way that enhanced colony success in a habitat-specific manner.

In our view, arguing about the precise semantics of “group selection” and “group adaptation” is beside the point: we should focus instead on the fascinating biology observed herein (Birch 2019). We also note that neglecting group selection when individual and colony level processes do not conflict can lead to inaccurate estimations on the nature of selection (Eldakar and Wilson 2011).

Three theoretical models have since attempted to resolve the issue of levels of selection in the *A. studiosus* system (Gardner 2015, Smallegange and Egas 2015, Biernaskie and Foster 2016). All models present elegant alternative explanations of the phenomena described in the original paper. However, these are phenomenological models, and thus ignore many empirical findings. For example, Biernaskie and Foster (2016) assumes that being aggressive is a “cheater” strategy, from which only the actor gains. This is not so (Pruitt and Riechert 2010, Wright et al. 2014). Smallegange and Egas (2015) assumes that the behavioural phenotype expression is strongly determined by the interaction of the environment and group size, in a heritable way, but not by parental phenotypes per se. Although this scenario is theoretically possible, no empirical evidence seems to support it (Pruitt and Riechert 2009a, Pruitt and Goodnight 2014, Purcell and Pruitt 2019). Finally, although Gardner (2015) demonstrated that the observed size-composition relationships can be described with an inclusive fitness approach, the assumptions of his model does not relate to the system’s known biology. We agree with Biernaskie and Foster (2016) that “*debates in sociobiology will benefit from stronger links between ecologically informed models and data*”. Yet, we would emphasize the part ecologically informed, more so than even their valuable model. Social spiders provide us with a unique chance to construct theoretical models where nearly all important assumptions are based on empirical data. Here we leverage these strengths to create a comprehensive individual-based model that integrates as much of the system’s known biology as possible.

Interestingly, there is but one element of the *Anelosimus studiosus* female life cycle of which nearly no data had been published, and that is the dispersal pattern of maturing individuals. Although originally dispersal was considered to weaken multilevel selection, in reality it can even facilitate it by generating a greater variability in group compositions (Eldakar and Wilson 2011). Clearly, the decision rules by which each female chooses between staying in the maternal web or leaving it, must be of crucial importance regarding the future composition of the focal colony. As both individual and colony fitness is driven by colony extinction events (Pruitt et al. 2017), the probabilities of which hinge on colony compositions (Pruitt and Goodnight 2014), we expect dispersal behaviour to be central from a multilevel selection point of view.

In the present study we aim to illuminate the importance of multilevel selection in structuring the dynamics of this system. By modeling both the individual- and colony-levels of a structured population we obtain a multilevel selection approach, hence we do not track relatedness or inclusive fitness (Kramer and Meunier 2016). We define within-colony selection as the evolution of traits based on the differential survival and reproduction of individuals. Similarly, following Wilson and Wilson (2007), we define between-colony selection as the evolution of traits based on the differential survival and reproduction of colonies.

Our study addresses three goals: 1) Building a detailed in silico model that is as strongly rooted in empirical data as possible. 2) Designing evolutionary simulations, where the dispersal behaviour of maturing females is the target of selection, and giving testable predictions regarding these. 3) Evaluating the effects and importance of multilevel selection in structuring this system.

## Methods

Our individual-based model contains a metapopulation of *A. studiosus* colonies, where we track each colony, and every individual in each colony, too. This is not a spatial model, neither on the level of the metapopulation, nor on the level of colonies. Competition between colonies is indirect: only a density-dependent selection is present for optimal nesting places over rivers (Jones and Riechert 2008, Little et al. 2019).

We consider two different environments in which colonies reside, the High Resource Environment (HRE) and the Low Resource Environment (LRE) (Pruitt and Goodnight 2014). There are two major differences between these environments, the colony extinction rules and the dispersal behaviour of individuals. In the following we will outline the model structure.

### Model process

At the beginning of the simulation we place 1000 solitary females into the metapopulation. As all adult females die at the end of the season (Jones et al. 2010), social colonies only appear whenever at least two offspring do not leave their maternal web. Individuals differ in their behavioural phenotypes (aggressive or docile), and colonies differ in their size and phenotypic composition (frequency of aggressive individuals).

**Figure 1:**
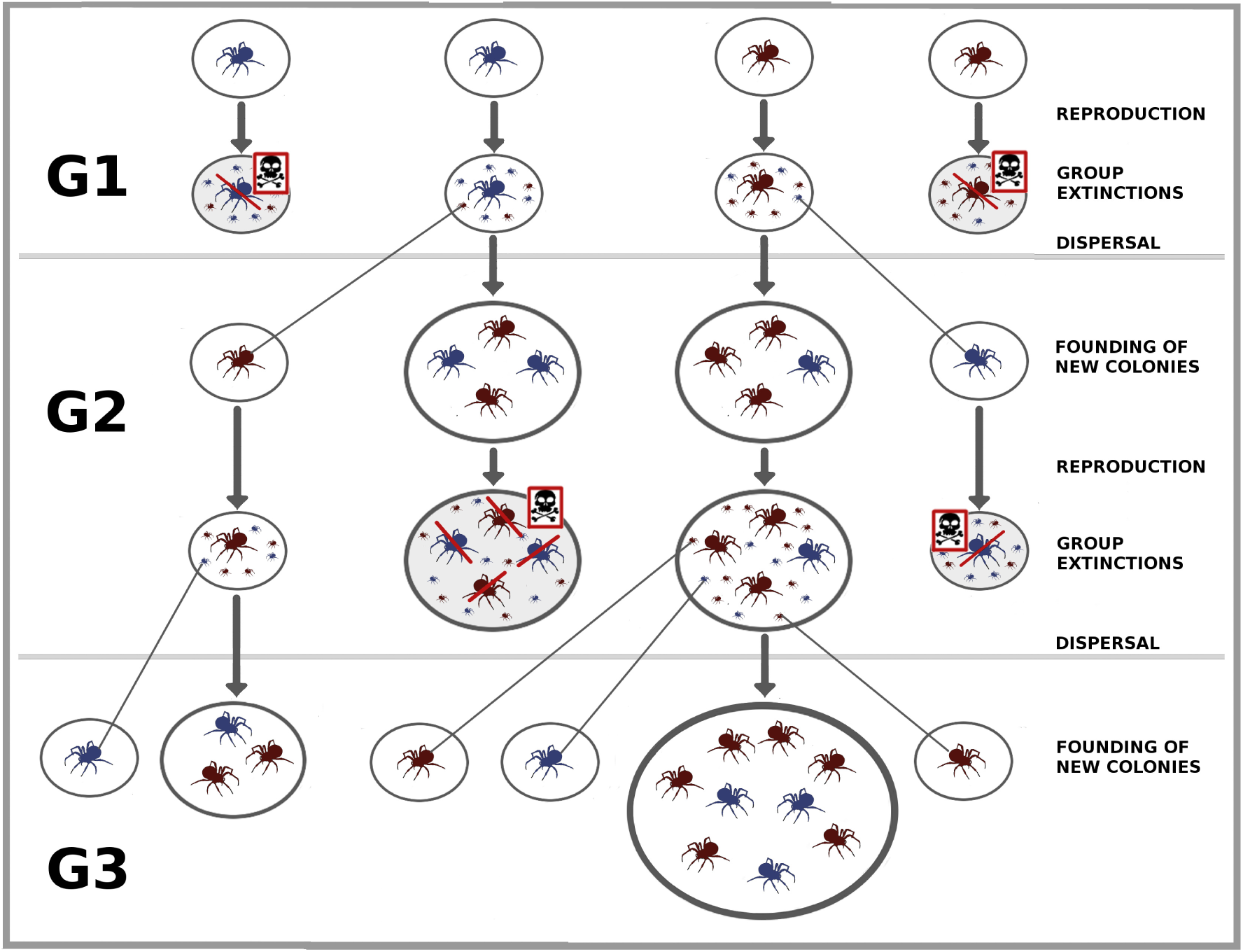
Schematic figure of the model process. Colonies are represented by the encircled spiders. The differing shades stand for the two phenotypes. The size of the colonies are proportional to the number of spiders within, with single spiders representing single-female colonies. Colonies with crossed spiders note extinction events. Vertical arrows denote time, grey lines denote dispersal events. Generations are separated by horizontal grey lines.

Each female reproduces once in her lifetime, thus we have non-overlapping generations. Before the instars reach adulthood, colonies have a probability of going extinct, depending on their size and phenotypic composition (Pruitt and Goodnight 2014). This process is the main contribution of between-colony selection in our model. In surviving colonies, maturing offspring either stay in their maternal web or leave it to found new single-female colonies. Dispersing individuals face the difficulty of finding a viable nesting place, that is, there is a density-dependent success or failure in founding a new colony.

The next generation, and all subsequent generations contain both the successfully founded new single-female colonies, and older multiple-female colonies. This way, although the generations of individuals are non-overlapping, the generations of colonies do overlap. In most cases we ran simulations for 1000 generations, and used the data of the last 500 generations for statistical analysis.

In the following we discuss all the above mentioned elements of the model briefly, but leave the mathematical details in the Appendix A. For the schematic figure of the model process see Figure 1.

### Food capture and reproduction

The amount of food a multiple-female colony captures depends both on the size and the composition of the colony (Pruitt and Riechert 2010, 2009a) Although the total food captured is a constantly growing function of the colony size, the per capita food intake peaks at 6-7 individuals, and decreases rapidly at larger colony sizes (Riechert and Jones 2008, Yip et al. 2008) Prey capture success increases with the proportion of aggressive individuals, but so does the cost of resource infighting (Pruitt and Riechert 2010, 2009a). Thus, based on the results of Lichtenstein and Pruitt (2015), we set the optimal colony composition with regard to food intake to be 20% aggressive individuals.

Mating is slightly disassortative (Pruitt and Riechert 2009a, Pruitt et al. 2011). We assume that the number of offspring is a linear function of the mother’s body mass, based on a common macro-trend (Marshall and Gittleman 1994). Aggressive females enjoy a benefit in reproductive potential over docile females (Jones et al. 2010, Pruitt and Riechert 2009a). This effect represents within-colony selection in the model.

Offspring phenotypes are determined by the parental phenotypes, and do not change throughout an individual’s lifetime (Pruitt and Goodnight 2014). In *A. studiosus*, generally there is a minor bias towards the representation of the aggressive phenotype, and no clutches are composed exclusively of either phenotype (Pruitt and Riechert 2009b, Pruitt and Goodnight 2014). Consequently, we determine phenotypes as follows: on average 90% of the offspring of two aggressive individuals are aggressive, 60% of the offspring of a mixed couple are aggressive, and 20% of the offspring of two dociles are aggressive. (For all the algorithmic and mathematical details visit Appendix A.)

### Colony extinction

In natural *A. studiosus* populations colony extinction events are the major determinants of both individual and colony fitness (Pruitt et al. 2017). Pruitt and Goodnight (2014) found that High and Low Resource Environments create characteristically different colony size and composition relationships. Thus, in our model the extinction probability of colonies depend on how far they lie from the optimum size-composition relationships obtained from their census data (see also on Figure 4, or more details on Figure A4). Consequently, the size-composition combination of colonies is the main target of between-colony selection.

In nature, single-female colonies face a larger risk of going extinct (Lichtenstein et al. 2018), as they only rely on the survival of one adult female, who bears the burden of all colony maintenance tasks. Thus, in the model, single-female colonies face an additional risk of going extinct. (For further details see Appendix A.)

### Instar survival, dispersal, density dependence

Maturing females either stay in the natal web or disperse to found new single-female colonies. As we had little information about the dispersal behaviour of this species, we designed evolutionary simulations to obtain adaptive dispersal curves for the model (see below). After the extinction and dispersal events all the adult females from the previous generation die (Jones et al. 2010).

### The evolution of the dispersal curves

We have conducted evolutionary simulations to obtain estimates about optimal dispersal behaviour in this system.

A growing colony size means reduced per capita food intake and increased local resource competition (Yip et al. 2008). Therefore we assumed the dispersal probability of a female to be an increasing function of her natal colony size. We decided to define this increasing function as a sigmoid curve, which can take very different forms depending on its parameter values. A dispersal curve is determined by three parameters: *α* defines the steepness of the inflexion point. *β* is the bottom limit of the sigmoid, that is, its minimum value, where (1 *− β*) is the upper limit of the same sigmoid. Finally, *γ* gives the position of the inflexion point (see Figure 2). (For the exact functions and further details see Appendix A.)

During the evolutionary simulations each offspring inherited the above described three parameters from her mother, but each parameter mutated with a certain probability, independently from each other. The mutant trait value was chosen from a normal distribution with an expected value of the parental trait (from the mother’s side). We ran simulations, where selection continuously shaped the form of the dispersal curves independently for the two phenotypes in the two environments. Thus, we followed the evolution of three parameters per phenotype per environment. We started the simulations with only solitary individuals, that is, with curves where instars always disperse. Simulations were run until the curves exhibited no more qualitative change, or for a maximum of 50,000 generations. As a consequence of the process we used, the final curves can be considered adaptive dispersal curves.

For all the basic parameter values used in the simulations, see Table A1.

## Results

Previously we specified our primary goal as the building of a detailed in silico model deeply rooted in empirical data. Yet, before discussing this objective, we present the outcome of the evolutionary simulations first, as the model builds upon these results.

### The evolution of the dispersal curves

During the evolutionary simulations we were looking for the adaptive value of the dispersal curve parameters in each of two environments. The top row of Figure 3 shows two typical cases of how the dispersal curves evolved in these simulations. (For an example of the parameter value evolutions, see Figure A6.) These observations suggest that the two phenotypes should behave differently in the two environments. In High Resource Environments, aggressive individuals typically remain in their natal webs, while docile individuals disperse, and the opposite is true in the Low Resource Environment (see also on Figure 3).

**Figure 2:**
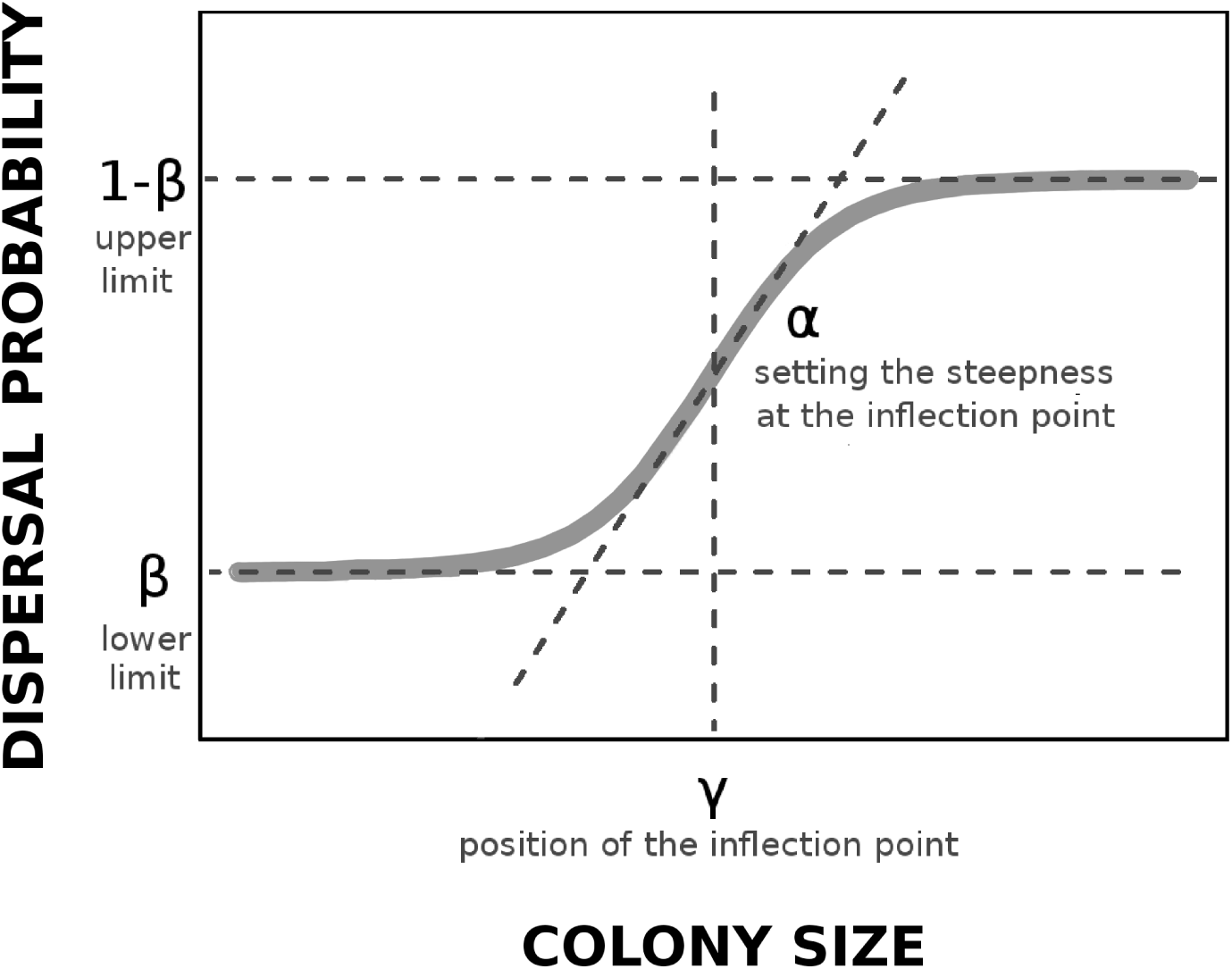
Sigmoid dispersal probability, depending on colony size, *n*_*i*_. The curve, *p*(*n*_*i*_) = *β* + (1 *−* 2*β*) (1 + *e^−α^*^(*ni−γ*)^ has three parameters: *α* sets the steepness of the inflexion point (the exact steepness here is (1 *−* 2*β*)*α/*4), *β* is the bottom limit or minimum value of the sigmoid, ((1 *− β*) is the upper limit), and *γ* is the position of the inflexion point.

The final curves were similar in the two environments for the opposing phenotypes (see the darkest curves in the top row of Figure 3 or the bottom row of Figure A6). Therefore, we decided to use the exact same curves in the two environments, only switching phenotypes across environments. According to unsystematic tests, these minor simplifications did not distort the behaviour of the model. For the final form of the dispersal curves, see the bottom line of Figure 3, and for the final parameters see Appendix A.

### Basic model outcomes

With the basic parameter set, which contains the empirically-based parameter values (see Table A1), and the dispersal curves obtained from the evolutionary simulations, the model closely recapitulated the size-composition relationships observed in nature. For a comparison of the simulated data and the census data of Pruitt and Goodnight (2014), see Figure 4. (We note here that only a qualitative comparison of the simulation and census data are possible, as the census data sampling was not random (J. N. Pruitt, personal communication).)

**Figure 3:**
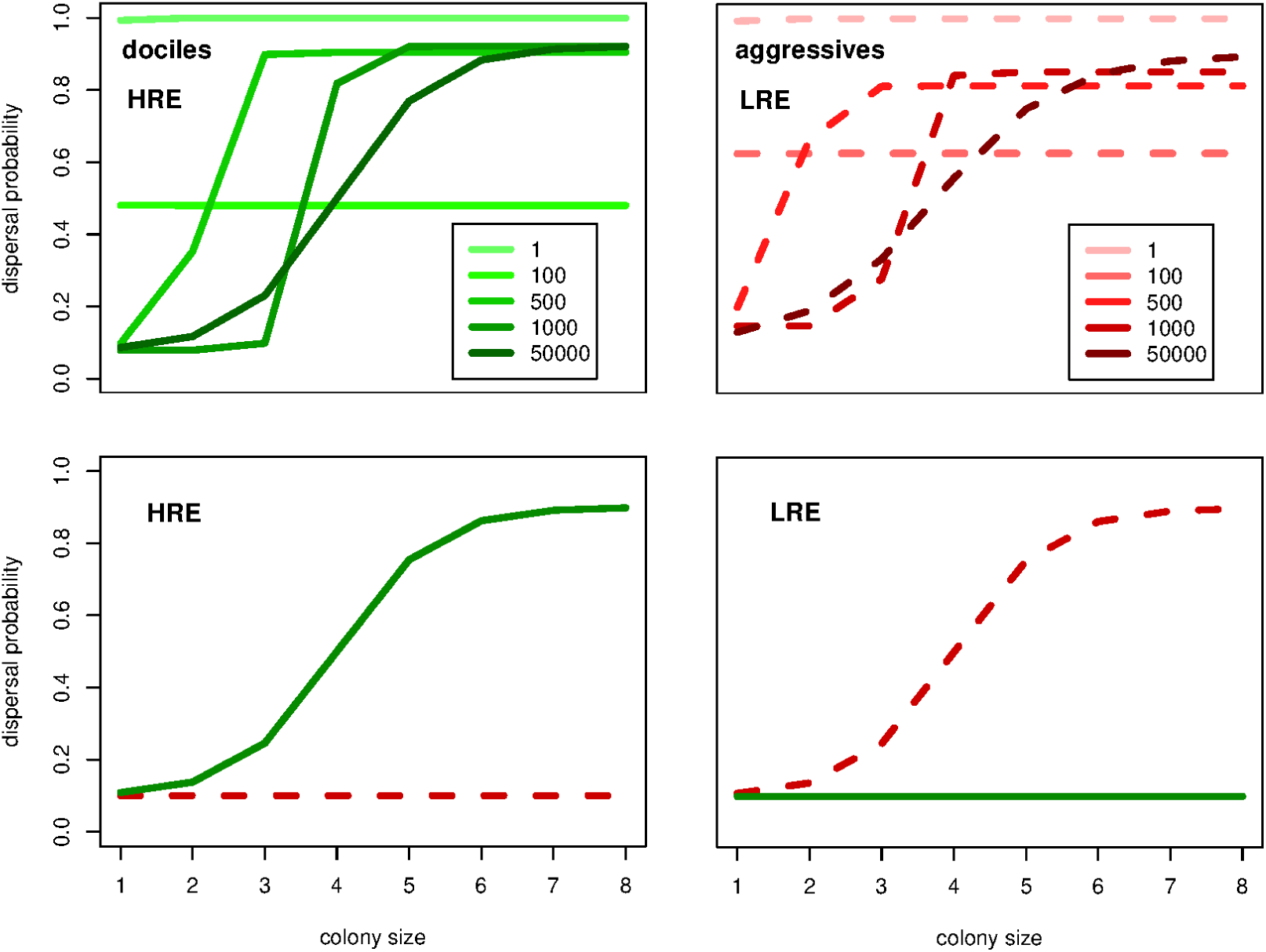
Evolution (top) and final form (bottom) of the dispersal probability curves. The red dashed lines represent the aggressive, and the green solid lines represent the docile phenotype. On the top plots curves are calculated from the given generation’s average values from generations 1, 100, 500, 1000 and 50 000. HRE stands for the High, LRE the Low Resource Environments. For other parameters see Table A1.

It is also notable that not only are the final colonies close to the optimal curves, but they are also quite diverse both in size and composition, just like colonies in natural populations. This is a somewhat unexpected result, as we did not set it an explicit goal throughout the evolutionary simulations, and because we dismissed stochasticity at numerous points from the model (e.g., neither the captured amount of food, or the number of eggs a given type of female laid had any variance).

**Figure 4:**
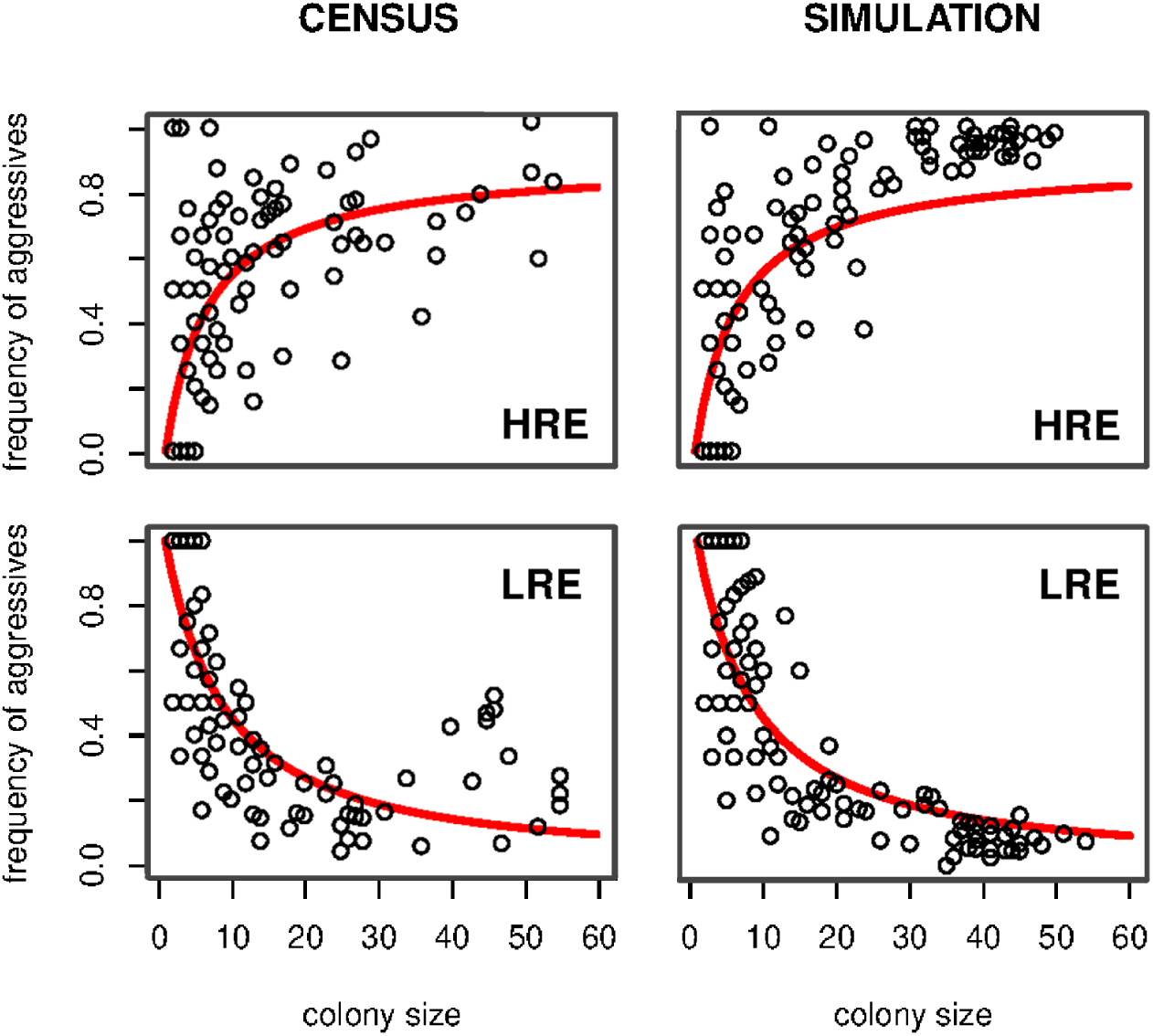
Frequency of aggressives plotted against colony size in the last generation with the adaptive dispersal probabilities (right) in comparison with the census data of Pruitt and Goodnight (2014) (left). For comparison, each plot cointains 150 randomly sampled data points from the respective datasets. Red lines are fitted curves of the census data of Pruitt and Goodnight (2014), used as colony size-composition optima during the simulations. HRE stands for the High, LRE the Low Resource Environments. For other parameters see Table A1 and Appendix A.

We have conducted simulations where there were no multifemale colonies, only single-female ones. During these tests, one of the phenotypes was always only marginally present. This is not surprising, as our extinction rules give a large benefit to one of the phenotypes in single-female colonies. In contrast, social populations with multiple-female colonies maintain a stable coexistence of phenotypes, since the optimal phenotypic composition of colonies changes rapidly with size in both environments. Consequently, if there are larger colonies present, the phenotype that is marginal in asocial population has to become more frequent.

### Robustness of the model

We have performed numerous robustness checks on the model. In the following we summarise the conclusions of these, but place most details in Appendix A. and figures in Appendix B.

#### Basic parameters

First, we have tested the robustness of the model against most parameters that were at least partly arbitrary. For instance, while we had empirical data confirming that single-female colonies face a higher extinction risk, we had to decide how high this additional risk should be in the basic model. Thus, we tested how the model reacted to varying extinction risk of single-female colonies. Based on these tests we concluded that the model was qualitatively robust to parameter variability within a reasonable range. (For further details on these parameter checks, see Table A1 and Table A2 in Appendix A).

#### Variance generating mechanisms

In *A. studiosus*, the differential survival of colonies is due to the diverse phenotypic composition of colonies. This diversity is the result of their mating system and the non-perfect heritability of the behavioural phenotypes. When we dismissed both variance generating mechanisms, one of the phenotypes completely disappeared from the populations (see Figure B3). Furthermore, the final colonies were very small, as homogeneous multiple-female colonies are generally less successful in both environments. The above results suggests that phenotypic variance generating mechanisms are necessary for a flourishing social population.

#### Within-colony selection

We have incorporated within-colony selection by defining unequal baseline fecundities for the different phenotypes. In the basic model, aggressive individuals have a 25% advantage in reproductive potential over docile individuals. Varying this advantage, or providing docile individuals with it did not alter the model outcomes qualitatively. Finally, the model did not change qualitatively even when we added within-colony negative frequency dependence (as described in Pruitt et al. (2017)), where in each colony the rarer phenotype enjoyed a reproductive advantage. This suggests that within-colony processes, though they may affect some demographic traits, are not essential for colony formation or for recapitulating the optimal size-composition relationships.

### Levels of selection and dispersal behaviour

Finally we have run a set of evolutionary simulations to test how the presence or absence of selection on each level affects the evolution of the dispersal curves. We have varied the within-colony selection as described above, and the between-colony selection by either incorporating or dismissing differential extinction events. That is, in one case used the previously defined colony extinction procedure (see section Colony extinction in Methods), and in the other assumed equal probability of extinction for all colonies. We have combined all the above settings in both environments.

Our results showed that if there was no between-colony selection, the size-composition distribution of colonies looked very similar in the two environments. It applied for all of these simulations that without between-colony selection, the population ends up similar to the Low Resource populations observed in nature (see also on Figure 5). The reason for this is that the model assumes an optimal aggressive frequency of 0.2 with regard to food capture success (Lichtenstein and Pruitt 2015). This composition is quite close to the optimal composition of large Low Resource colonies. As the foraging efficiency of colonies decreases rapidly with size, large colonies with suboptimal composition have very low reproductive output. That is why selection in our model drives aggressive individuals to leave their natal web with a high probability. It is also noteworthy that when we varied the direction of within-colony selection, we have seen no qualitative differences between the resulting size-composition distributions during these evolutionary simulations. Within-colony selection merely affected how large colonies typically got before collapsing, and thus affected the mean colony sizes. For an example see Figure 5, where we present the size-composition distributions with varying between-colony selection and the basic parameter set.

These results support the finding of Pruitt et al. (2017) that although within-colony selection can clearly affect the colonies’ size and composition, the characteristic patterns of the different environments are driven by site-specific colony extinction events.

**Figure 5:**
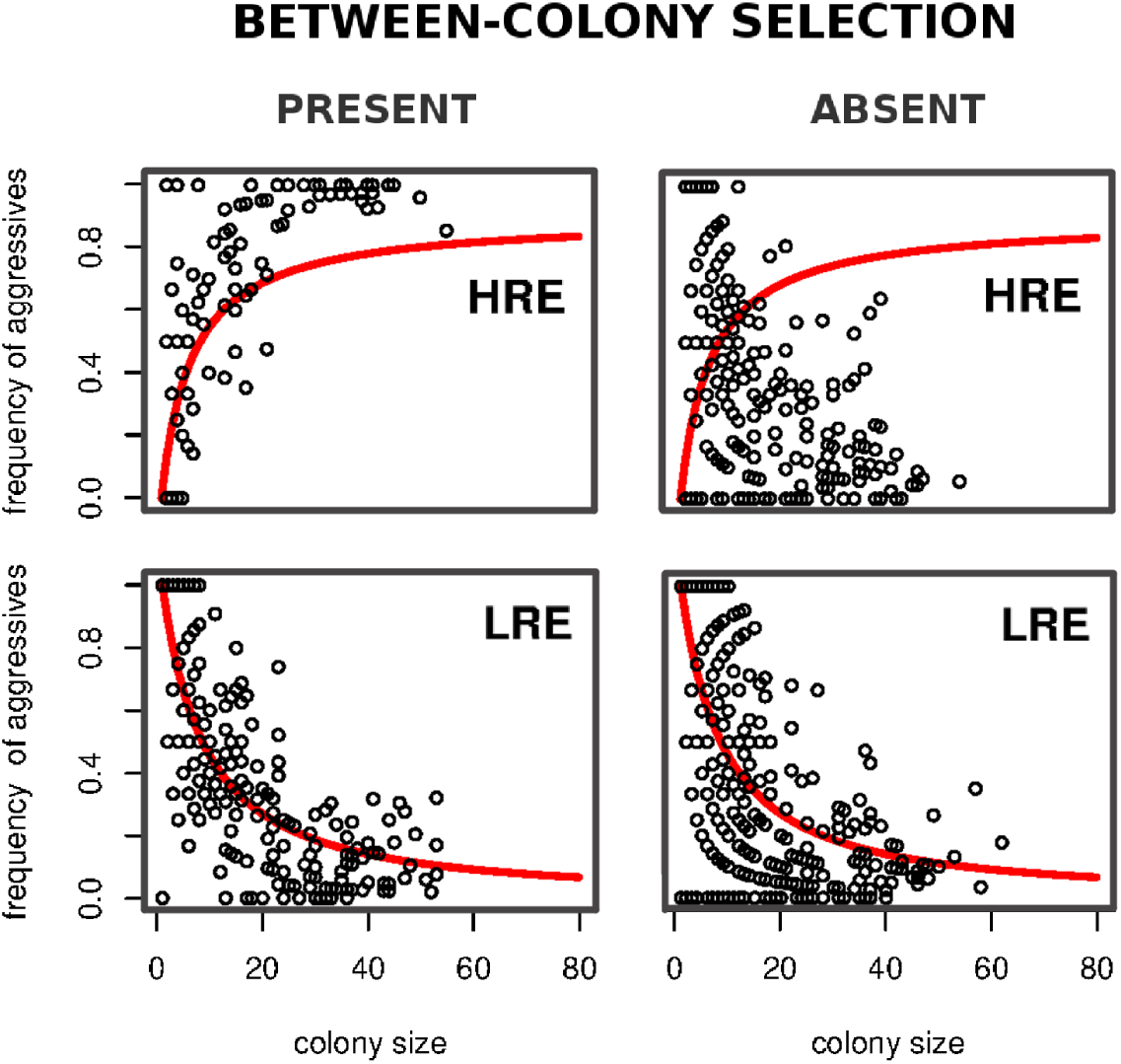
Frequency of aggressives plotted against colony size in the last generations of evolutionary simulations. The left column shows populations with between-colony selection and the right column shows populations without between-colony selection. HRE stands for the High, LRE the Low Resource Environments. Red lines are fitted curves of the census data of Pruitt and Goodnight (2014), used as colony size-composition optima during the simulations. For other parameters see Table A1.

## Discussion

The spider *A. studiosus* offers an exceptional opportunity to observe and understand how multilevel selection acts in nature. So far, only empirical studies and phenomenologocal models have been deployed to deepen our understanding of this species’ dynamics. Here we developed a detailed individual-based model that incorporates most of the known properties of this system. With adaptive dispersal curves, obtained in evolutionary simulations, we observed a close resemblance between the demographics of our in silico populations and those seen in situ.

We found that the Low Resource Environment of the model generated greater population sizes and enhanced population stability. This either means that with a given number of optimal nesting places Low Resource populations indeed can reach higher density, or that these populations face yet hidden additional difficulties. According to the model simulations, such difficulty could be an increased instar mortality risk or higher risk for single-female colonies, both of which could be investigated experimentally.

Our model demonstrates that populations should evolve phenotypic-biased dispersal in opposing direction, and that this behaviour alone can coarsely generate the size-composition characteristic of High vs. Low Resource sites. The results are in agreement with the finding of Pruitt (2013) that in Low Resource Environments, daughter colonies founded by docile individuals were significantly closer to the mother colony than colonies founded by aggressive individuals, if we regard this distance as a proxy for dispersal propensity.

We tested the model with no differential extinction events, where the only difference between the two environments were the opposing, but not evolvable dispersal curves. Surprisingly, we obtained similar demographic outcomes as with the basic model (see Figure B2)). This suggest that the typical size-composition distributions in the two environments are highly dependent on the different dispersal behaviour of individuals. Pruitt and Goodnight (2014) found that experimental colonies with suboptimal size-composition combinations typically collapse. However, such colonies would not be present in natural populations at all, if naturally occurring *A. studiosus* exhibited the dispersal behaviour they evolve in models. It is the prospect of future studies to describe the dispersal behaviour of subadult *A. studiosus*, and thus to evaluate the power of dispersal behaviour in natural populations. Nevertheless, this model has clearly shown that evolving an environmentally driven dispersal behaviour can be a straightforward way to produce optimal group compositions.

The dispersal pattern of individuals can help to determine their colonies’ success, but this behaviour also affects the fitness of the dispersing individual. Differential phenotypic-biased dispersal is an adaptation that only exists in the context of a social population, as a population composed of only subsocial individuals exhibits an all-disperse rule (Furey 1998). Thus, it could be contended that dispersal could be a kind of colony level adaptation that enhances the performance of the colony and which only exists in the context of social populations. However, this view remains open to the criticisms that the decision to disperse is a behaviour of an individual that reflects its relative odds of performing well in ailing colony versus a pioneering foundress.

Multi-female colony forming in *A. studiosus* increases with decreasing temperature (Furey 1998, Jones et al. 2007, Riechert and Jones 2008). We have found here that increasing the extinction risk of single-female colonies generate a situation where sociality is needed for population persistence (see Figure B1). Our model is consistent with the findings of Jones et al. (2007), where colder environments increased extinction risk of single-female colonies, thus favoring multifemale colonies. Although Jones et al. (2007) concentrated on the prolonged development of spiderlings and the associated increased risk of orphaned offspring in cold climates, their data as easily fit a model of increased mortality risk from any source in cold environments. Our model is more agnostic than Jones et al. (2007) by modeling any generic risk of increased mortality in singleton nests. Consistent with our model, small colonies of *A. studiosus* experience far lower survivorship irrespective of sites’ temperature (Lichtenstein et al. 2018). Although this model is not designed for answering the question of how sociality evolved in this species, it is still notable that we detected a critical parameter range where multifemale colony forming is required for populations to persist. Furthermore, we also found that the formation of multifemale colonies alters phenotype frequencies in the population, and thus stably maintains the coexistence of the different behavioural types.

Multifemale *A. studiosus* colonies are more than just safety measures against too early colony extinctions. Multifemale colonies can capture larger prey (Furey 1998, Ward 1986), and perform an effective division of labour (Wright et al. 2014). Such emergent qualities, cooperation and division of labour, give the strong impression of a higher level of organisation, and a possible unit of selection. Selection on such a higher level of organisation is only possible if there is variation, and differential survival or reproduction between the units.

In *Anelosimus studiosus* there is a large variation between colonies with regard to their size and composition. This variation is partly generated by the mating system and the non-perfect heritability of the behavioural phenotypes. But, according to our results, the variation is also generated by the site and phenotypic biased dispersal of maturing females. As the differential extinction of colonies is the major driver of fitness in this species, the second prerequisite of colony-level selection, the differential survival or reproduction of colonies, is also fulfilled. If dispersal behaviour indeed is a major influence on colony composition, then it must be the target of not just individual-, but also colony-level selection. Dispersal behaviour is thus both the outcome, and trigger of multilevel selection in this social spider

## Declarations

Funding for ZsV and ISch was provided by the National Scientific Research Fund (OTKA K128289) and by Hungary’s Economic Development and Innovation Operative Program (GINOP 2.3.2-15-2016-00057). Funding for JNP was provided by the Canada 150 Research Chair Program

ZsV implemented the simulations and drew the figures. All three authors contributed to the study design and the manuscript.

