## Appendix for "Dispersal behaviour as the outcome and trigger of multilevel selection in a social spider"

### Appendix A: Methods and Results details

#### Food capture

One generation of individuals lives for $t = 100$ days. Daily foraging success of a multiple-female colony depends on its size ($n_i$) and composition ($q_i$, the frequency of aggressives) in the following way:

$$C_i^{(m)}(n_i, q_i) = F(n_i)\ B_{\mu_c,w_c}(q_i) \tag{A1}$$

$$F(n_i) = \frac{n_i^2 + a}{n_i^2 + b}$$

where $C_i^{(m)}$ is the expected amount of prey the $i^{th}$ multiple-female colony captures, $F(n_i)$ is the prey capture success as a function of the colony size and $B_{\mu_c,w_c}(q_i)$ is a scaled bell shaped curve, determined by it expected value, $\mu_c$ and the parameter setting its width, $w_c$ (Lichtenstein and Pruitt 2015). Here, and from now on, $B_{\mu,w} = exp\big(-(\frac{1}{2}(x-\mu)/w)^2\big)$. This curve has a maximum value of 1 at its expected value. (For $F(n_i)$ and for $B_{\mu_c,w_c}$ see Figure A1 and Figure A2, respectively.) $a$ and $b$ are constants scaling the $F(n_i)$ function.

For single-female colonies the expected daily food intake is

$$C_i^{(s)}(1) = 0.9\ F(1) = 0.9\ \frac{1+a}{1+b}.$$

Thus, the expected per capita food capture will be $\frac{1}{n_i}\ C_i^{(m)}(n_i, q_i)$ (see Figure A1). For the sake of simplicity we assume that the weight gain of individuals is a linear function of the consumed energy. That is, $\frac{t}{n_i}\ C_i^{(m)}(n_i, q_i)$ determines the condition of females in the $i^{th}$ colony.

Aggressives probably both consume and use more energy (Lichtenstein and Pruitt 2015), thus within colonies we assume roughly equal gains. The expected amount of food colonies get has no variance here. Consequently, colony members always enjoy the same physical condition.

#### Mating

At the end of a generation's time mating takes place. We find a random mate slightly disassortatively for every female independently from the fellow colony members' mates (Pruitt and

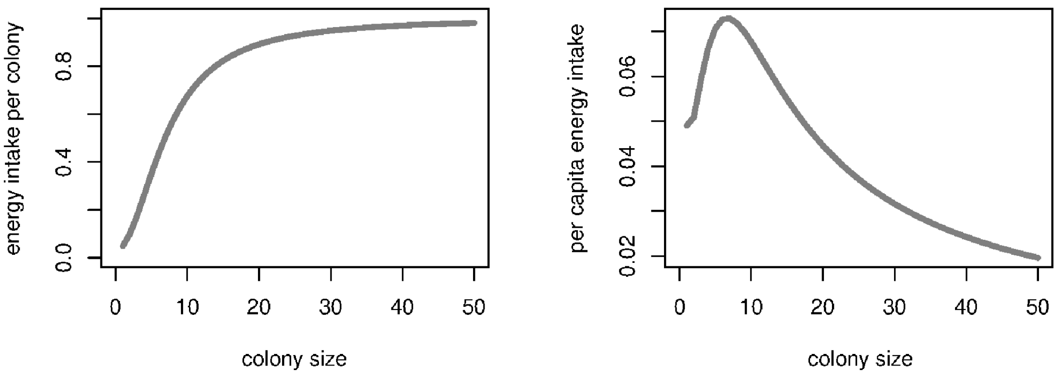


**Figure A1: Colony (left) and per capita (right) food capture success, $F(n_i)$, as a function of colony size.** $a = 1.5$, $b = 50$.

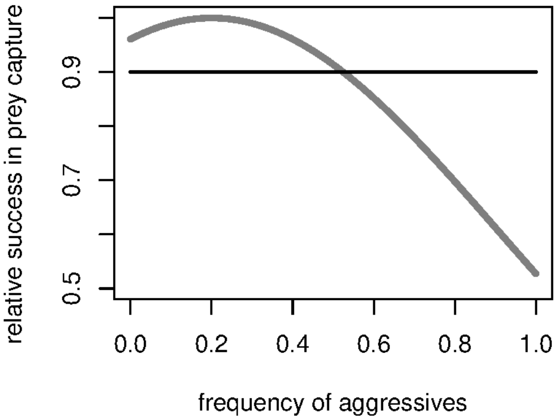


**Figure A2: Colony food capture success, $B_{\mu_c,w_c}(q_i)$, as a function of colony composition.** $\mu_c = 0.2$, $w_c = 0.5$. **The grey line represents multiple-female colonies, the black line represents single-female colonies.**

Riechert 2009a, Pruitt et al. 2011). Both phenotypes have a 55% chance of having a mate with the opposite phenotype. Every female produces an eggsac. In our model, aggressives are more fecund than dociles. Although there is evidence that this benefit stands only in single colonies (Jones et al. 2010), as the probability for the two phenotypes having exactly equal fecundity is practically zero, we have chosen aggressives to enjoy this advantage in all types of colonies. In the model single aggressive females produce $o_A = 20$, single dociles produce $o_D = 16$ eggs. (From now on, the indices $A$ and $D$ denote aggressives and dociles, respectively.) The size of an eggsac is a linear function of the mother's body mass (Marshall and Gittleman 1994). We scale the "cost" of an egg so as single aggressive females to have 20 eggs, that is, to be $(\frac{1}{20}\ T\ C_i^{(s)}(1))$. Thus, the number of eggs in a multiple-female colony for aggressives and dociles, respectively, are

$$
\begin{aligned}
o_A &= int\Big(\frac{t\ C_i^{(m)}(n_i, q_i)}{\frac{1}{20}\ t\ C_i^{(s)}(1)}\Big) = int\Big(20\ \frac{C_i^{(m)}(n_i, q_i)}{C_i^{(s)}(1)}\Big) \\
o_D &= int\Big(1.25\ o_A\Big) \qquad \text{(A2)}
\end{aligned}
$$

where $int\Big( * \Big)$ rounds down its argument to the nearest integer. During the evolutionary simulations the number of eggs is chosen from a Gaussian distribution with $\mu_A = o_A$, $\mu_D = o_D$ and $\sigma = 5$ (see later).

In one model version the expected number of eggs was multiplied by a negatively frequency dependent coefficient, $\phi$: the more frequent a female's phenotype in the colony, the less eggs she produces, and vice versa (Pruitt et al. 2017), according to the following equations:

$$
\begin{aligned}
\phi_A &= (1.5 - \epsilon - (1 - 2\epsilon)\ q_i) \\
\phi_D &= (1.5 - \epsilon - (1 - 2\epsilon)\ (1 - q_i)) \qquad \text{(A3)}
\end{aligned}
$$

for aggressives and for dociles, respectively, where $\epsilon \in (0, 0.5)$ defines the strength of frequecy dependence (see Figure A3). Thus the expected number of eggs were $o'_A = o_A\ \phi_A$ and $o'_D = o_D\ \phi_D$. In the frequency dependent model version $\epsilon = 0.3$.

##### Colony extinction and density dependence

The optimal colony composition for every colony size varies with the environment. The optimal composition is given by a fitted curve, for which we have used the census data of Pruitt and Goodnight (2014). we used a general two parameters saturation function for the High, and a general two parameters convex decreasing function for the Low Resource Environment.

The fitted curves are as follows:

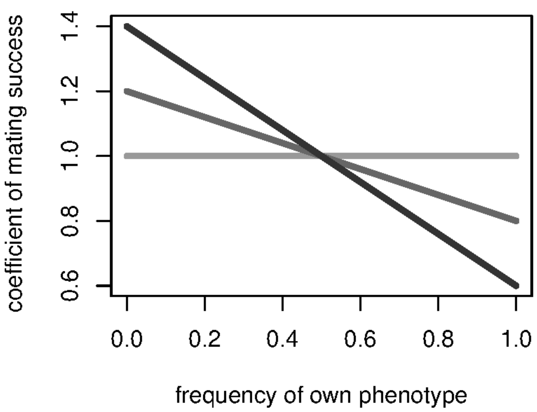


**Figure A3: The negative frequency dependent coefficient of the number of eggs. From lighter to darker grays $\epsilon = 0.5$ (no frequency dependence), $\epsilon = 0.3$ and $\epsilon = 0.1$.**

$$\frac{h_1\ n}{h_2 + n} - \frac{h_1}{h_2 + 1}$$

$$\left(\frac{l_1 + n}{l_1 + 1}\right)^{-l_2}$$

for the High and the Low Resource Environments, respectively, where $n$ is the colony size and the fitted parameters are $h_1 = 1.0830$, $h_2 = 4.8451$, $l_1 = 9.33015$ and $l_2 = 1.25641$ (see also on figure A4).

The extinction probability ($p_{ext}(n_i, q_i)$) we define based on the colony's distance from the optimum composition and is given by an inverted and scaled bell shaped curve which never reaches 1 and always exceeds 0 (see the top plot of Figure A4), as follows:

$$p_{ext}(n_i, q_i) = 0.95 - B_{0,w_{ext}}(d_i)/\ \lambda, \tag{A4}$$

where $d$ is the colony's distance from the optimal composition, $w_{ext}$ defines the strength of composition based selection and $\lambda$ scales the baseline extinction risk for all colonies. In the basic model $\lambda = 2.0$ and $w_{ext} = 0.25$. Single-female colonies face an additional 20% risk of going extinct.

In surviving colonies, all instars have a constant probability of survival independent of their phenotype. Dispersing individuals face a density-dependent success in founding a new colony,

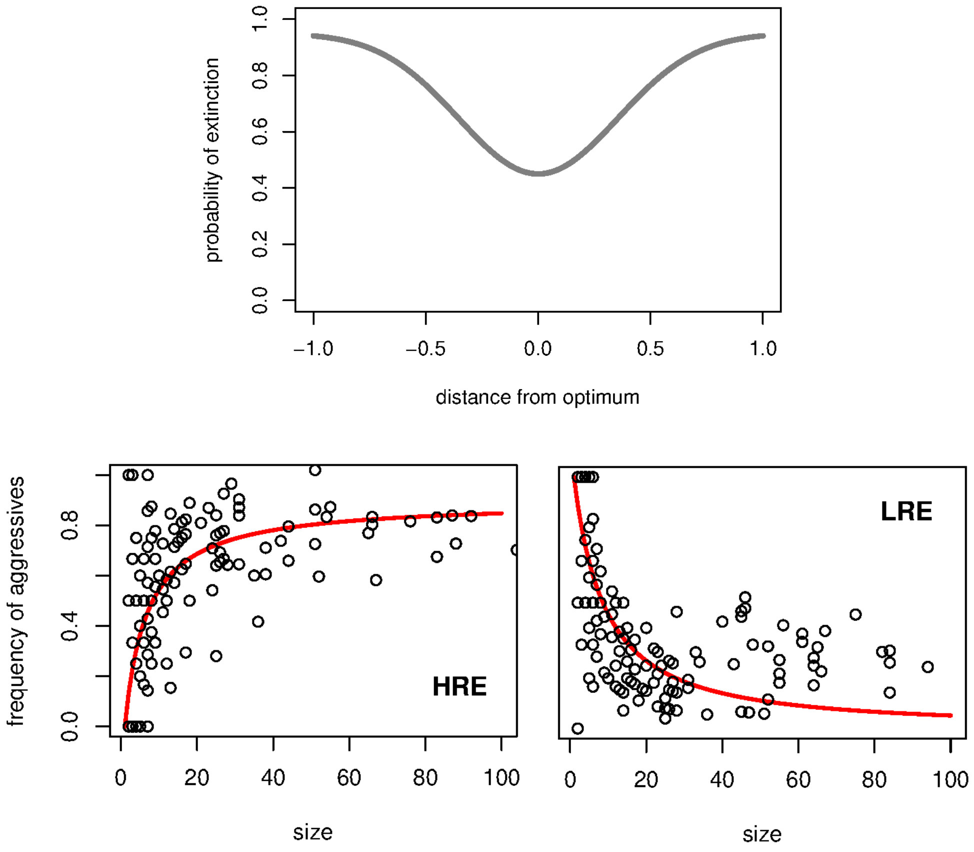


**Figure A4: Extinction probability ($p_{ext}$) given by the colony's distance from the optimal composition (top) and optimal compositions in two environments for varying colony sizes. Data points represent the census data from Pruitt and Goodnight (2014). HRE stands for the High, LRE the Low Resource Environments.**

because there are a finite number of nesting places. The probability of success is given by

$$\frac{\delta}{C + D + \delta}, \tag{A5}$$

where $\delta$ scales the population size, $C$ is the number of currently occupied nesting places including collapsed colonies, and $D$ is the number of dispersing individuals (see Figure A5). In the basic model $\delta = 1000$.

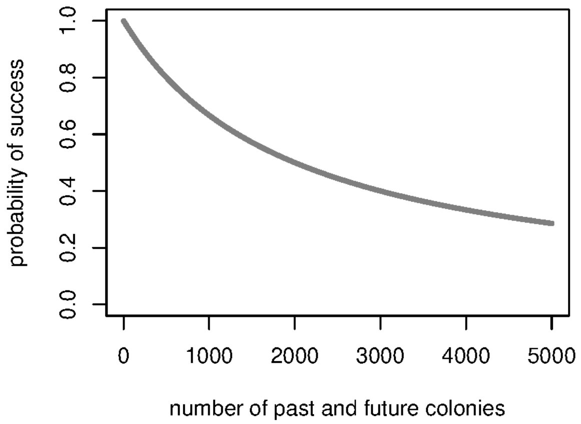


**Figure A5: Densitiy dependent success in founding a new colony. The $x$ axis shows the sum of occupied nesting places and dispersing females ($(C + D)$ from Equation A5). $\delta$=1000.**

##### The evolution of the dispersal curves

We were looking for the optimal dispersal probability functions $p_A(n_i)$ and $p_D(n_i)$ in the two environments. To be more precise, we were looking for the three parameters defining a sigmoid curve: $\alpha$ sets the steepness of the inflexion point (the exact steepness here is $(1 - 2\beta)\alpha/4$), $\beta$, the bottom limit of the sigmoid, that is, its minimum value, where $(1 - \beta)$ is the upper limit of the same sigmoid, and finally, $\gamma$, the position of the inflexion point.

Formally:

$$p_A(n_i) = \beta_A + \frac{1 - 2\beta_A}{1 + exp(-\alpha_A(n_i - \gamma_A))} \tag{A6}$$

$$p_D(n_i) = \beta_D + \frac{1 - 2\beta_D}{1 + exp(-\alpha_D(n_i - \gamma_D))} \tag{A7}$$

for aggressives and dociles, respectively. The initial parameter combinations resulted in practically all individuals leaving their natal web ($\alpha_{A,0} = \alpha_{D,0} = 5$, $\beta_{A,0} = \beta_{D,0} = 0$, $\delta_{A,0} = \delta_{D,0} = 0$). During the evolutionary simulations each offspring inherited the above described three parameters from her mother, but each parameter mutated with a certain probability, independently from each other. The mutant trait value was chosen from a normal distribution with an expected value of the parental trait (from the mother's side). The larger the mutation probability and the variance of the normal distribution, the faster the evolution. During these simulations the mutation probability was 0.1 and the mutant trait value's normal distribution had a standard deviation of 0.1.

Figure A6 shows that some parameters (e.g. $\alpha_D$ in the High or $\alpha_A$ in the Low Resource Environment) soon reach some sort of equilibrium value, while others (e.g. $\alpha_A$ in the High or $\alpha_D$ in the Low Resource Environment) seems to make a random walk suggesting neutral selection in the given range. If we look at the evolution of the curves themselves (see Figure 3), it gets more obvious while such wanders of the parameters can happen. The optimal dispersal probability curve of the "staying" phenotype in both environments is a linear line around 0.1. As there is no visible inflexion point, its steepness and position is relatively irrelevant, as long as the curve roughly has this linear shape at the proper value. Correspondingly, the parameter values of the "leaving" phenotypes in both environments all reach equilibrium values, and indeed, the dispersal curves of these phenotypes show a characteristic and similar shape.

We have run several simulations with various parameter combinations, and the dispersal curves became quite similar in all of these unsystematic tests. Thus we have decided to choose such round numbers for these parameters, which are close to these equilibrium values, and which can be used for both environments (but obviously for the opposing phenotypes). As a result of the evolutionary simulations thus we have chosen the parameters $\alpha_A = 5$, $\alpha_D = 1.5$, $\beta_A = 0.9$, $\beta_D = 0.1$, $\gamma_A = -2$, and $\gamma_D = 4$ for the High Resource Environment, and $\alpha = 5$, $\alpha_A = 1.5$, $\beta_D = 0.9$, $\beta_A = 0.1$, $\gamma_D = -2$, and $\gamma_A = 4$. for the Low Resource Environment.

For the final curves, determined by the above parameters, see Figure 3.

#### Parameters checks

Table A1 contains the basic parameter set and their range in which the model were tested. In Table A2 we have summarised the trends in reaction to parameter variation.

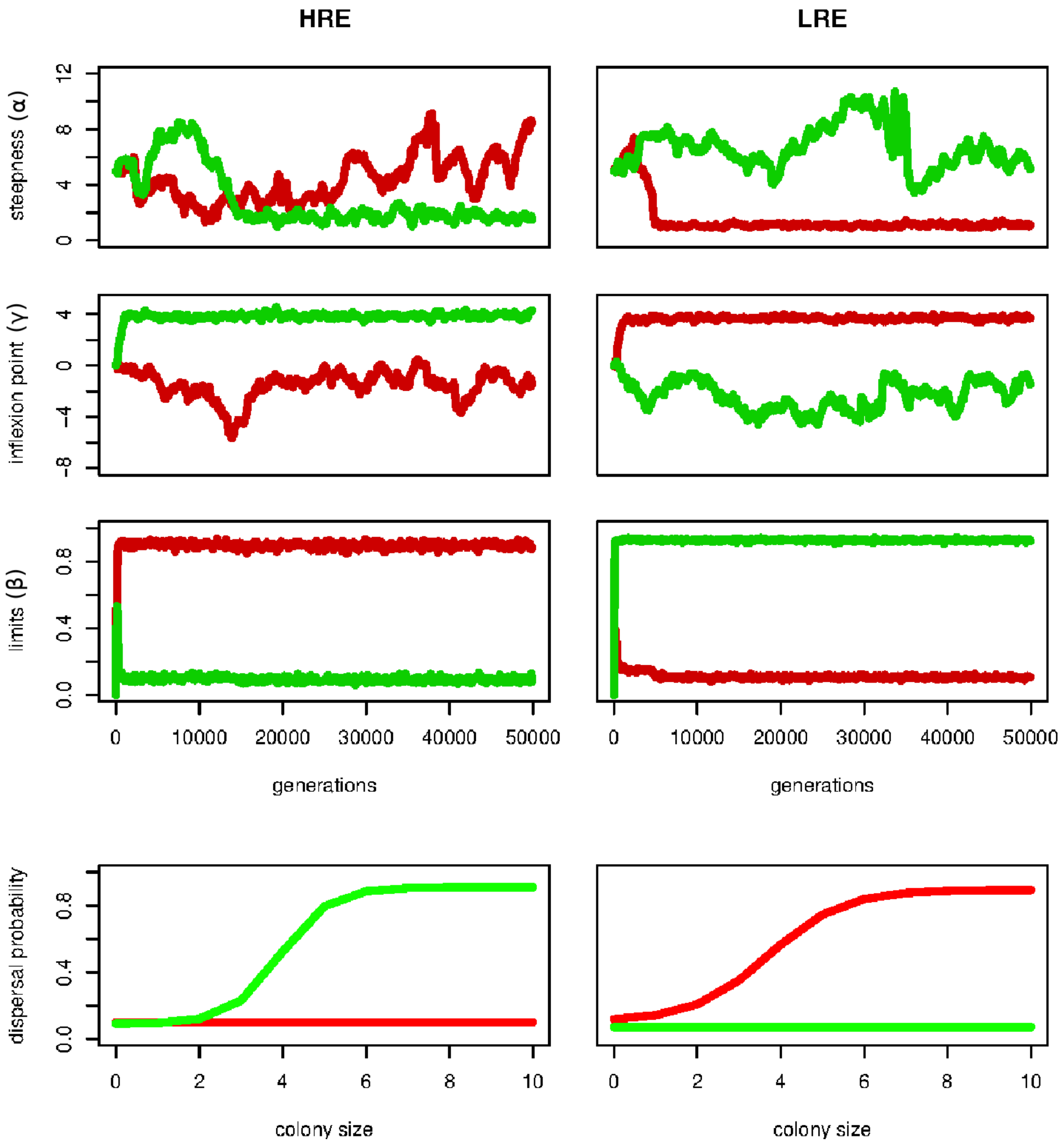


**Figure A6: Evolving parameters in the two environments. The left colums shows the High Resource Environment (HRE), the right shows the Low Resource Environment (LRE). Red lines represents parameters of the aggressive phenotype, green lines those of the docile phenotye. The first row (steepness) shows the parameter $\alpha$ (the exact steepness here is $(1-2\beta)\alpha/4$), the second (inflexion point) the parameter $\gamma$, and the third (minimum and maximum) shows the parameter $\beta$. The bottom plots show the dispersal curves defined by mean values of the last 10 000 generations.**

| | notion | basic value | range tested |
|---|---|---|---|
| relative foraging success of single-female colonies | | 0.9 | 0.5 – 1.3 |
| strength of frequency dependence | $\epsilon$ | 0.5 | 0.1 – 0.5 |
| baseline fecundity (A/D) | | 20/16 | 11 – 25 |
| parameter scaling the baseline extinction risk | $\lambda$ | 2.0 | 1.25 – 2.5 |
| additional extinction risk of single-female colonies | | 0.2 | 0.0 – 0.6 |
| instar mortality | | 0.75 | 0.65 – 0.85 |
| a parameter scaling the density dependence | $\delta$ | 1000 | 500 – 2500 |

**Table A1: Parameter names, notions (if applicable), their value in the basic parameter set, and their range in which the model were tested.**

| | mean # of colonies | mean colony size | % of singles |
|---|---|---|---|
| basic (HRE) | 459 | 6.84 | 0.59 |
| basic (LRE) | 993 | 7.5 | 0.54 |
| higher foraging success of singles | ↑ | ↑ | ↓ |
| higher fecundity of either types | ↑ | ↑ | ↓ |
| lower extinction risk for all colonies | ↑ | ↑ | ↓ |
| lower single additional extinction | ↑ | ⊘ | ⊘ |
| weaker frequency dependent fecundity | ↑ | ↑ | ↓ |
| lower instar mortality | ↑ | ↑ | ↓ |
| weaker density dependence | ↑ | ⊘ | ⊘ |

**Table A2: End-of-generation demographic data of the basic setting and parameter dependent trends. Arrows represent statistically significant correlational trends in both environments with $p < 0.05$ in all cases, and $0.01 < p < 0.05$ in only such cases where the relationship was clearly nonlinear. Basic refers to the basic model, where values are calculated from several runs' data.**

### Appendix B: Supplementary Figures (Results)

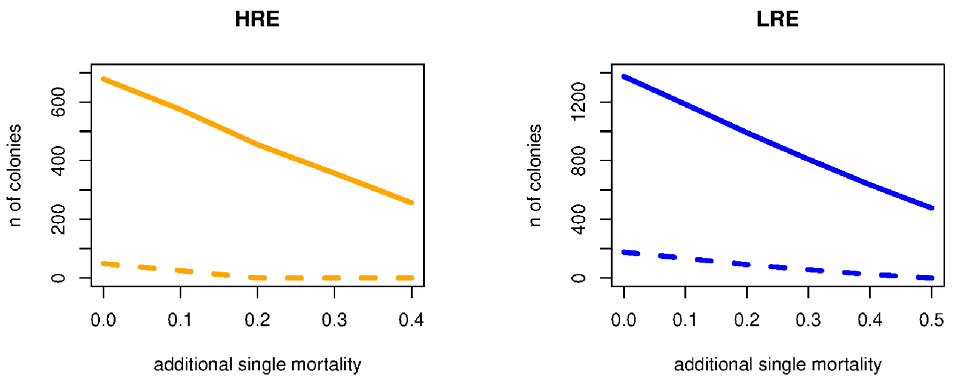


**Figure B1: Population size with varying additional mortality of single-female colonies in both environments, in both social and asocial populations. The orange lines represent the High Resource Environment (HRE, left), the blue lines the Low Resource Environment (LRE, right). The dashed lines represent the asocial, the solid lines the social populations.**

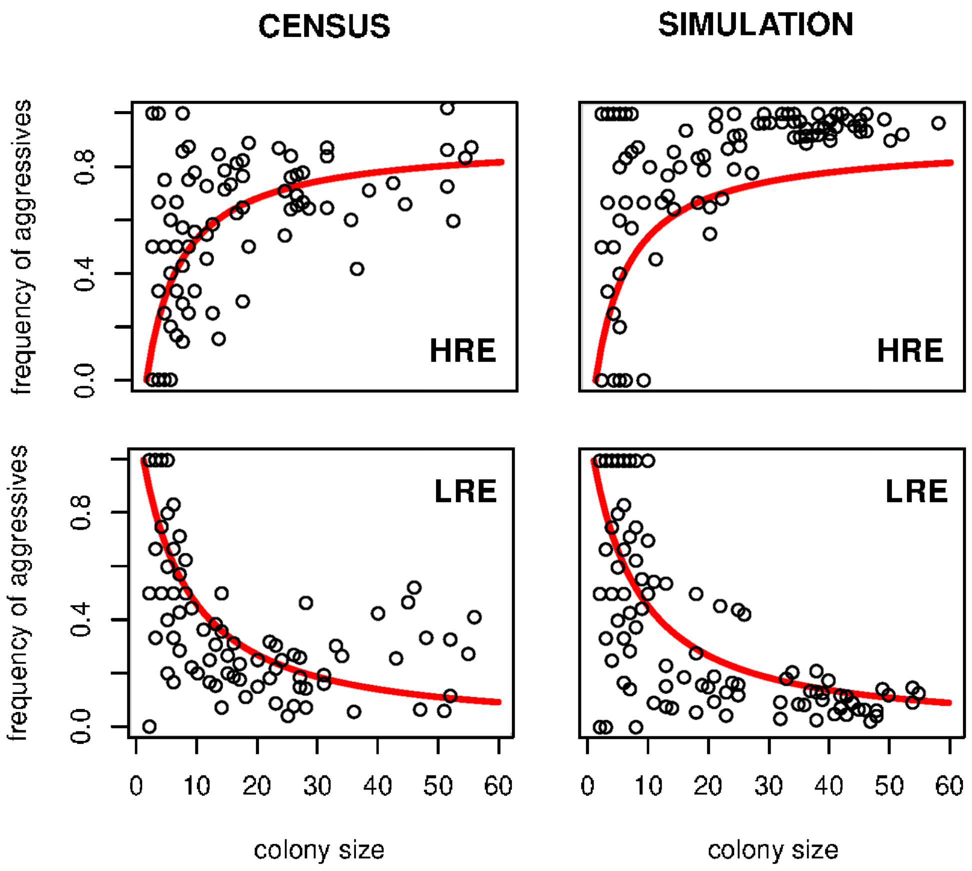


**Figure B2:** Frequency of aggressives plotted against colony size in the last generation with no environment driven extinction rule (right) in comparison with the census data of Pruitt and Goodnight (2014) (left). For comparison, each plot cointains 150 randomly sampled data points from the respective datasets. Red lines are fitted curves of the census data of Pruitt and Goodnight (2014). HRE stands for the High, LRE the Low Resource Environments. In this simulation the only difference between the environments is the dispersal pattern of individuals.

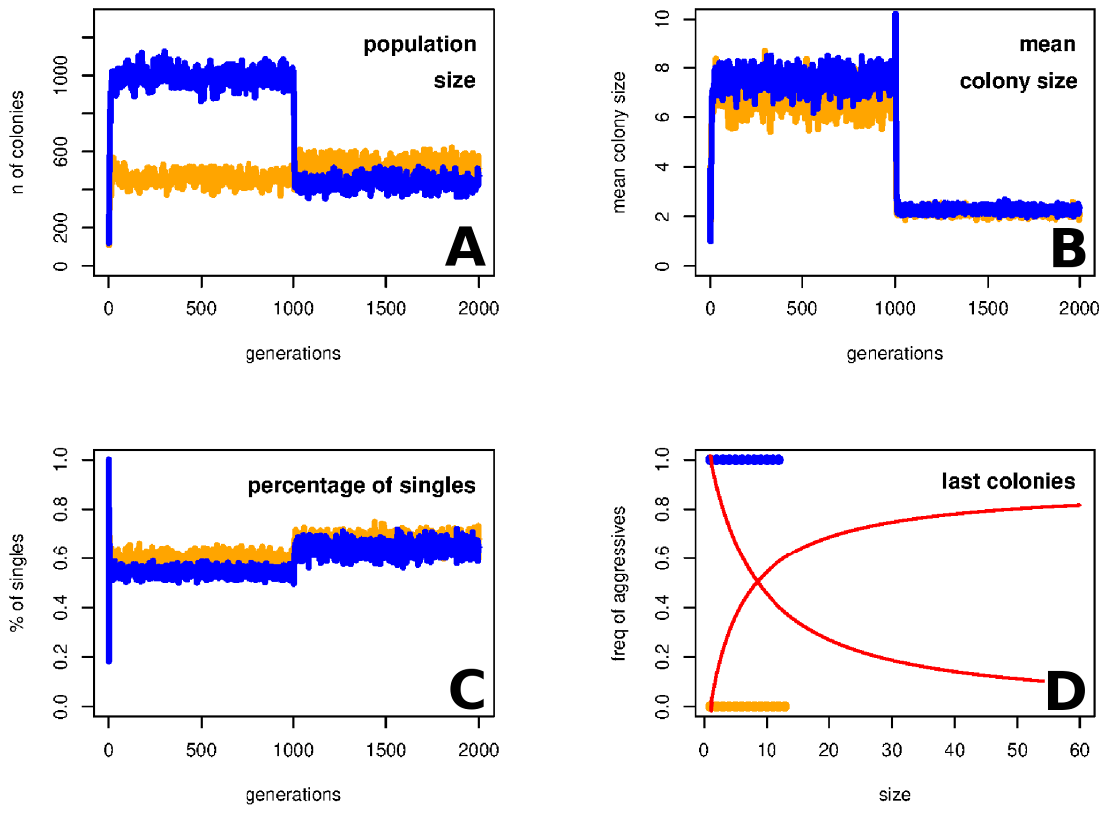


**Figure B3: Demographic data and the final populations in the model version, where after 1000 generations we have cut all variance generating mechanisms (disassortative mate choice and imperfect inheritance). Orange represents the High Resource Environment and blue represents the Low Resource Environment. Plot A shows the number of colonies throughout the simulation. Plot B shows the mean colony sizes within generations. Plot C shows the percentage of single-female colonies from all colonies. Plot D shows the size and composition of the last generation of the simulations in both environments. The logic of this plot is the same as the logic of Figure 4: the solid red lines represent the optimal colony size-composition relationships (from the census data of Pruitt and Goodnight (2014)). The two environments are pictured together, because only homogeneous docile or aggressive colonies are present, thus the data points of the two environmnets do not hide each other.**
